# Xpert MTB/XDR: A ten-color reflex assay suitable for point of care settings to detect isoniazid-, fluoroquinolone-, and second line injectable drug-resistance directly from *Mycobacterium tuberculosis* positive sputum

**DOI:** 10.1101/2020.09.08.288787

**Authors:** Yuan Cao, Heta Parmar, Rajiv L. Gaur, Deanna Lieu, Shobana Raghunath, Nova Via, Simone Battagalia, Daniela M. Cirillo, Claudia Denkinger, Sophia Georghiou, Robert Kwiatkowski, David Persing, David Alland, Soumitesh Chakravorty

**Author notes:** Division of Tropical Medicine, Center of Infectious Diseases, University Hospital of Heidelberg, Germany (Claudia Denkinger). These authors contributed equally to this study (Yuan Cao, Heta Parmar, David Alland, Soumitesh Chakravorty). Author order was determined on the basis of seniority.

## Abstract

We describe the design, development, analytical performance and a limited clinical evaluation of the 10-color Xpert MTB/XDR assay (CE-IVD only, not for sale in the US). This assay is intended as a reflex test to detect resistance to Isoniazid (INH), Fluoroquinolones (FLQ), Ethionamide (ETH) and Second Line Injectable Drugs Drugs (SLID) on unprocessed sputum samples and concentrated sputum sediments which are positive for *Mycobacterium tuberculosis*. The Xpert MTB/XDR assay simultaneously amplifies eight genes and promoter regions in *M. tuberculosis* and analyzes melting temperatures (Tms) using sloppy molecular beacon probes (SMB) to identify mutations associated with INH, FLQ, ETH and SLID resistance. Results can be obtained under 90 minutes and requires 10-color GeneXpert modules. The assay can differentiate low versus high-level resistance to INH and FLQ as well as cross-resistance versus individual resistance to SLIDs by identifying mutation-specific Tms or Tm patterns generated by the SMB probes. The assay has a Limit of Detection comparable to the Xpert MTB/RIF assay and succesfully detected 16 clinically significant mutations in a challenge set of clinical isolate DNA. In a clinical study performed at two sites with 100 sputum and 214 clinical isolates, the assay showed a sensitivity of 94-100% and a specificity of 100% for all drugs except for ETH when compared to sequencing. The sensitivity and specificity when compared to phenotypic drug susceptibility testing were in the same range. Used in combination with a primary tuberculosis diagnostic test, this assay is expected to expand the capacity for detection of drug-resistant tuberculosis.

## Introduction

Drug-resistant *Mycobacterium tuberculosis* remains a significant threat to global Tuberculosis (TB) care and public health. The World Health Organization (WHO) has estimated that in 2017, 3.4% of new cases and 18% of previously treated cases had rifampicin-resistant TB (RR-TB) or multidrug-resistant TB (MDR-TB), and as many as 8.5% of new TB cases had extensively drug-resistant TB (XDR-TB)(1). MDR-TB is caused by *M. tuberculosis* that is resistant to Isoniazid (INH) and Rifampicin (RIF), and XDR-TB is additionally resistant to at least one of the Fluoroquinolones (FLQ) and Second Line Injectable Drugs (SLIDs) including Amikacin (AMK), Capreomycin (CAP), and Kanamycin (KAN). Phenotypic drug susceptibility testing (P-DST), the current gold standard for identifying drug resistance in *M. tuberculosis*, takes 6 to 8 weeks to provide definitive results and poses a bio-hazard risk for laboratory personnel, especially when working with XDR strains. Thus, treatment is often empirically based on other factors such as past medical history or local prevalence of resistance (2, 3). Delays in appropriate treatment can increase both mortality and transmission of drug resistant strains.

The Xpert MTB/RIF (*in vitro* diagnostic use only) assay and its more advanced version, the Xpert MTB/RIF Ultra (CE-IVD only; not for sale in the US) assay (Cepheid, CA, USA) were designed to simultaneously detect the presence of *M. tuberculosis* and RIF-resistance in an integrated and fully automated system directly from sputum. These assays can be used with only minimal training at point of care settings, (4, 5) where widespread implementation has led to an overall decrease in TB incidence in some studies (6) and increased notifications of RIF-resistance in others (7). However, these assays only detect RIF-resistance, and it has been shown that selection of TB treatment regimens based only on detection of RIF resistance can result in suboptimal therapy for 49% of patients with MDR or XDR TB (8). Thus, additional tests that identify resistance to INH, FLQs and SLIDs equally rapidly in similar point of care settings are also necessary. The MTBDR*plus* and MTBDR*sl* version 2.0 (HAIN Life Sciences, Germany) assays are currently recommended by WHO as molecular tests to detect INH and RIF, and FLQ and SLID resistance, respectively. However, these assays are not suitable for near patient or point of care testing, as it requires a sophisticated laboratory settings (9).

We had previously developed and validated a prototype cartridge-based assay that detected *M. tuberculosis* resistant to INH, the FLQs and SLID directly from sputum using a multiplexed 10 color molecular test. This cartridge was a proof of principle initial prototype, which underscored our technical capability to develop a functional 10 color reflex test on sputum samples positive for *M. tuberculosis* by primary diagnostic tests, including the Xpert MTB/RIF and the Xpert MTB/RIF Ultra assay. The cartridge detected resistance with high sensitivity and specificity directly from the sputum of TB patients in less than 2 hours (10, 11). However, since the development of the prototype cartridge, mutations in *fabG1* and *oxyR-ahpC* intergenic regions have also been shown to account for INH resistance (12) (13); mutations at the −14 position in the *eis* promoter region have been shown to confer resistance to both KAN and AMK (other *eis* promoter mutations only confer resistance to KAN) (14, 15); while the mutation “a1401g” in *rrs* gene confers cross resistance to AMK, KAN and CAP (16, 17); and mutations in the quinolone resistance-determining region (QRDR) in *gyrA* have been subdivided into mutations that cause low-level resistance (*gyrA* A90V, *gyrA* S91P, and *gyrA* D94A) and other mutations that elevate MICs to later-generation of FLQ drugs more substantially (18-20). The MIC differences caused by these two different categories of mutations have also been shown to have different clinical outcomes when a FLQ is used as part of MDR/XDR therapy (21, 22). In the current study, we describe a new, more advanced test, the Xpert MTB/XDR assay (CE-IVD only, not for sale in the US), that was redesigned to improve mutation coverage for INH, differentiate low level INH resistance from higher level resistance, identify Ethionamide (ETH) resistance, distinguish between low and high levels of resistance to FLQs, and identify cross resistance versus individual resistance to the SLIDs. We have also improved the overall sensitivity of the assay and reduced the time to result to less than 90 minutes after the run is started. Here we describe the development and initial clinical evaluation of this assay along with the additional attributes to improve assay performance.

## Results

### Detection and differentiation of *gyrA* QRDR mutations

We redesigned our previous assay in order to be able to specifically identify the *gyrA* QRDR mutations A90V, S91P and D94A that are associated with low level FLQ resistance (19, 23), and to distinguish them from the other QRDR mutations that are associated with higher level resistance. We designed three overlapping Sloppy molecular beacon (SMB) probes with slightly varying sequences against the *gyrA* QRDR. These three *gyrA* probes were designed to generate specific “three Tm window” patterns, which identify and distinguish each of the above mentioned QRDR mutations when they occur in the absence of other *gyrA* QRDR mutations. We designated one wild type (WT) window and multiple mutant windows for each of the *gyrA* probes. The gyrA1 and gyrA3 probes were assigned three mutant windows (MutA, MutB and MutC), and the gyrA2 probe two mutant windows (MutA and MutB). The multiple mutant and the single WT Tm for each probe can theoretically generate nearly 48 different “three Tm window” combinations specific to the QRDR mutations and the WT sequence. The WT QRDR sequence generates a “WT-WT-WT” Tm window pattern for the three probes and the mutant sequences have one or more of the WT Tm values replaced by mutant Tm (Table 1). Thus, the three overlapping probes with different binding affinities to the *gyrA* QRDR enabled us to generate tri-window Tm patterns specific to each of A90V, S91P and D94A mutations, namely “MutB-MutA-MutB”; “MutB-MutA-MutC” and “MutA-MutA-WT” respectively for the “gyrA1-gyrA2-gyrA3” probes as shown in Table 1. Each pattern is specific for one of these three mutations, enabling the identification of low FLQ resistance. Additionally, we designed the probes to be agnostic to the Ser/Thr polymorphism in the codon 95, so that identical Tm patterns for the WT and all the mutant sequences are generated for both these polymorphisms. We tested *gyrA* QRDR containing plasmids bearing 11 different *gyrA* mutation types for both the 95S and 95T polymorphisms as a challenge set as shown in Table 1. The probes successfully identified and differentiated the individual low resistance-conferring mutations from other mutations by their different Tm signatures (falling in different Tm windows), as indicated in Table 1.

**Table 1.** Mean Tm values (± SD)** of the three *gyrA* probes with representative *gyrA* mutant plasmids and their corresponding windows. WT windows are highlighted in green and the three mutant windows are in blue, orange and yellow respectively.

| Genotype |  | Probe 1 | Probe 2 | Probe 3 | FLQ Result Output |
| --- | --- | --- | --- | --- | --- |
| **WT**<br>**(95S or T)** | Mean T<sub>m</sub> | 76.2 (±0.2) | 70 (±0.2) | 70.8 (+0.2) | **FLQ Resistance NOT**<br>**DETECTED** |
|  | T<sub>m</sub> window | WT | WT | WT |  |
| **G88C**<br>**(95S or T)** | Mean T<sub>m</sub> | 72.7 (+0.3) | 65.2 (±0.4) | 66.4 (+0.3) | **FLQ Resistance**<br>**DETECTED** |
| | ΔT<sub>m</sub> | -3.4 | -4.8 | -4.4 | |
|  | T<sub>m</sub> window | Mut B | Mut B | Mut C |  |
| **G88A**<br>**(95S or T)** | Mean T<sub>m</sub> | 71.7 (+0.5) | 63.9 (±0.3) | 65.4 (+0.3) | **FLQ Resistance**<br>**DETECTED** |
| | ΔT<sub>m</sub> | -4.4 | -6.1 | -5.4 | |
|  | T<sub>m</sub> window | Mut B | Mut B | Mut C |  |
| **A90V**<br>**(95S or T)** | Mean T<sub>m</sub> | 72.2 (±0.2) | 75.6 (+0.3) | 76.2 (+0.2) | **Low FLQ Resistance**<br>**DETECTED** |
| | ΔT<sub>m</sub> | -4.1 | 5.6 | 5.4 | |
|  | T<sub>m</sub> window | Mut B | Mut A | Mut B |  |
| **S91P**<br>**(95S or T)** | Mean T<sub>m</sub> | 72.2 (±0.1) | 74.8 (±0.1) | 66.1 (±0.4) | **Low FLQ Resistance**<br>**DETECTED** |
| | ΔT<sub>m</sub> | -3.9 | 4.8 | -4.7 | |
|  | T<sub>m</sub> window | Mut B | Mut A | Mut C |  |
| **D94A**<br>**(95S or T)** | Mean T<sub>m</sub> | 78.9 (±0.2) | 73.4 (±0.2) | 71.4 (±0.1) | **Low FLQ Resistance**<br>**DETECTED** |
| | ΔT<sub>m</sub> | 2.7 | 3.4 | 0.6 | |
|  | T<sub>m</sub> window | Mut A | Mut A | WT |  |
| **D94G**<br>**(95S or T)** | Mean T<sub>m</sub> | 76 (±0.2) | 69.5 (±0.3) | 75.8 (±0.2) | **FLQ Resistance**<br>**DETECTED** |
| | ΔT<sub>m</sub> | -0.1 | -0.5 | 5 | |
|  | T<sub>m</sub> window | WT | WT | Mut B |  |
| **D94N**<br>**(95S or T)** | Mean T<sub>m</sub> | 72.9 (±0.3) | 66.1 (+0.4) | 68.9 (+0.3) | **FLQ Resistance**<br>**DETECTED** |
| | ΔT<sub>m</sub> | -3.2 | -3.9 | -1.9 | |
|  | T<sub>m</sub> window | Mut B | Mut B | WT |  |
| **D94Y**<br>**(95S or T)** | Mean T<sub>m</sub> | 72.5 (+0.3) | 65.1 (+0.4) | 68.6 (+0.3) | **FLQ Resistance**<br>**DETECTED** |
| | ΔT<sub>m</sub> | -3.6 | -4.9 | -2.2 | |
|  | T<sub>m</sub> window | Mut B | Mut B | Mut C |  |
| **D94H**<br>**(95S or T)** | Mean T<sub>m</sub> | 73.2 (+0.3) | 65.6 (+0.3) | 68.9 (+0.3) | **FLQ Resistance**<br>**DETECTED** |
| | ΔT<sub>m</sub> | -2.9 | -4.4 | -1.9 | |
|  | T<sub>m</sub> window | WT | Mut B | WT |  |
| **A90V+S91P** | Mean T<sub>m</sub> | 67.5 (+0.3) | 79.3 (+0.2) | 71.7 (+0.1) | **FLQ Resistance** |
| (95S or T) | ΔTm | -8.6 | 9.3 | 0.9 | DETECTED |
|  | Tm window | Mut C | Mut A | WT |  |
| A90V+G88C<br>(95S or T) | Mean Tm | 67.8 (±0.5) | 71.3 (+0.2) | 72.2 (+0.2) | FLQ Resistance<br>DETECTED |
|  | ΔTm | -8.3 | 1.3 | 1.4 |  |
|  | Tm window | Mut C | WT | WT |  |

\*\*Mean Tm and SD were generated from 3 to 163 replicates from multiple experiments depending on genotype

### Mutant panel challenge

We assessed the ability of the Xpert MTB/XDR assay to detect drug resistance associated mutations in clinical *M. tuberculosis* isolates. A panel of 14 clinical isolates with canonical mutations in the target genes and promoter regions known to be associated with clinical INH, ETH, FLQ, and SLID resistance were tested (Table 2). All the mutations were confirmed by P-DST and Sanger sequencing of each of the target genes and promoter regions. The assay was able to detect all the mutations (except a single *gyrB* mutation) and correctly determine the specific drug resistance profile for all the isolates tested. The assay correctly detected low-INH and ETH resistance conferring mutation (*inhA* c-15t) and all the other INH resistance-conferring mutations in *katG* (S315T), *fabG1 (*g603a) and *ahpC* (g-48a, g-6a). The assay also detected all the *gyrA* QRDR mutations, including a triple and a double mutant (Table 2). The assay resulted in correct “low FLQ resistance detected” calls for A90V, S91P and D94A mutations that are associated with low-level FLQ resistance and resulted in the correct “FLQ resistance detected” call for D94G and D94Y mutations. Isolates with mutations associated with SLID resistance in *eis* (g-10a, c-12t) and *rrs* (a1410g) genes were also identified correctly. Although the assay was able to correctly determine the resistance profile of all 14 isolates, it was not able to identify the *gyrB* mutation T539N because the Tm difference (dTm) between the WT Tm and this mutation was 1.1°C, which did not result in the Tm falling in the mutant window for *gyrB* probe. However, as the isolate also had an A90V mutation, it was identified as a low FLQ resistant sample. The *gyrB* T539N mutation has been reported to be present at a very low frequency in FLQ resistant isolates (24) and functional genetic studies have demonstrated that introduction of this mutation into the wild type *M. tuberculosis* genome does not result in any appreciable increase in MIC to third-generation FLQ (25). Thus, the assay’s failure to detect *gyrB* T539N would not be expected to affect sensitivity for detecting FLQ resistance. The specific 10 Tm profiles corresponding to each mutation, which also enables specific identification of each genotype associated with the target genes, are shown in Table 2.

**Table 2.** Xpert MTB/XDR mutant DNA panel challenge.

| Strain ID | Gene conferring resistance | Resistance conferring variant** | Melting temperature (± SD) ($^{\circ}$C) | | | | | | | | | |
| --- | --- | --- | --- | --- | --- | --- | --- | --- | --- | --- | --- | --- |
|  |  |  | inhA | katG | fabG | ahpC | gyrA1 | gyrA2 | gyrA3 | gyrB2 | rrs | eis |
| H37Rv | NA | None | 76.3<br>(+0.0) | 73.7<br>(+0.0) | 71.4<br>(+0.1) | 69.3<br>(+0.0) | 76.1<br>(+0.1) | 70.3<br>(+0.1) | 71.1<br>(+0.1) | 69.6<br>(+0.0) | 75<br>(+0.0) | 68.5<br>(+0.1) |
| Clinical isolate | katG | S315T | 76.5<br>(±0.0) | 68.4<br>(± 0.0) | 71.6<br>(±0.0) | 0<br>(±0.0) | 76.5<br>(+0.0) | 70.4<br>(+0.0) | 71.3<br>(+0.0) | 69.7<br>(±0.1) | 75.2<br>(+0.0) | 68.6<br>(+0.0) |
| Clinical isolate | gyrA | D94G | 76.5<br>(±0.1) | 68.4<br>(±0.1) | 71.5<br>(+0.0) | 0.0 | 76.2<br>(±0.1) | 69.7<br>(+0.0) | 75.7<br>(+0.0) | 69.7<br>(+0.1) | 71.0<br>(+0.0) | 68.6<br>(±0.1) |
|  | katG | S315T |  |  |  |  |  |  |  |  |  |  |
|  | rrs | a1410g |  |  |  |  |  |  |  |  |  |  |
| Clinical isolate | gyrB | C539A | 71.1<br>(+0.0) | 73.8<br>(+0.0) | 71.6<br>(+0.0) | 67.2<br>(+0.0) | 72.3<br>(+0.0) | 75.9<br>(±0.1) | 76.5<br>(+0.0) | 68.6<br>(+0.0) | 71.1<br>(+0.0) | 68.6<br>(+0.0) |
|  | rrs | a1410g |  |  |  |  |  |  |  |  |  |  |
|  | gyrA | A90V |  |  |  |  |  |  |  |  |  |  |
|  | OxyR-ahpC | g -6 a |  |  |  |  |  |  |  |  |  |  |
|  | inhA promoter | c -15 t |  |  |  |  |  |  |  |  |  |  |
| Clinical isolate | gyrA | D94Y | 76.3<br>(+0.0) | 68.2<br>(+0.0) | 75.6<br>(+0.0) | 0.0 | 72.7<br>(+0.0) | 0.0 | 69<br>(+0.0) | 69.6<br>(+0.0) | 75<br>(+0.0) | 68.5<br>(+0.0) |
|  | katG | S315T |  |  |  |  |  |  |  |  |  |  |
|  | fabG1 | g609a |  |  |  |  |  |  |  |  |  |  |
| Clinical isolate | gyrA | S91P | 71.1<br>(+0.0) | 73.9<br>(+0.1) | 71.7<br>(±0.1) | 0.0 | 72.2<br>(+0.0) | 75.1<br>(+0.0) | 66.6<br>(+0.0) | 69.7<br>(+0.0) | 75.2<br>(+0.0) | 68.6<br>(+0.0) |
|  | inhA promoter | c -15 t |  |  |  |  |  |  |  |  |  |  |
| Clinical isolate | gyrA | A90 A/V,<br>D94 D/G | 71.03<br>(±0.1) | 73.7<br>(+0.1) | 71.6<br>(+0.0) | 0.0 | 76.13<br>(±0.1) | 69.4<br>(±0.2) | 76<br>(±0.10) | 69.7<br>(+0.0) | 75.1<br>(+0.0) | 68.5<br>(+0.0) |
|  | inhA promoter | c -15 t |  |  |  |  |  |  |  |  |  |  |
|  | OxyR-ahpC | g -6 a |  |  |  |  |  |  |  |  |  |  |
| Clinical isolate | gyrA | G88G/A,<br>A90V,<br>S91S/P | 76.43<br>(±0.1) | 68.3<br>(±0.1) | 71.5<br>(±0.1) | 69.3<br>(+0.0) | 72.2<br>(±0.1) /<br>67.6<br>(±0.1) | 70.5<br>(+0.0)/<br>79.4<br>(+0.0) | 71.6<br>(+0.0) /<br>76.7<br>(+0.0) | 69.7<br>(±0.1) | 71<br>(±0.1) | 68.6<br>(±0.1) |
|  | katG | S315T |  |  |  |  |  |  |  |  |  |  |
|  | rrs | a1410g |  |  |  |  |  |  |  |  |  |  |
| Clinical isolate | katG | S315T | 76.4<br>(±0.1) | 68.3<br>(+0.0) | 71.5<br>(+0.0) | 67.1<br>(+0.0) | 76.4<br>(+0.0) | 70.3<br>(+0.0) | 71.3<br>(+0.0) | 69.6<br>(+0.0) | 75.1<br>(+0.0) | 68.6<br>(+0.0) |
|  | OxyR-ahpC | a -48 g |  |  |  |  |  |  |  |  |  |  |
| Clinical isolate | katG | S315T | 76.4<br>(±0.1) | 68.3<br>(±0.1) | 71.5<br>(±0.1) | 66.03<br>(±0.1) | 76.3<br>(+0.0) | 70.3<br>(+0.0) | 71.2<br>(+0.0) | 69.6<br>(±0.1) | 75.03<br>(±0.1) | 68.6<br>(±0.1) |
|  | OxyR-ahpC | a -48 g |  |  |  |  |  |  |  |  |  |  |
| Clinical isolate | gyrA | D94A | 70.9<br>(+0.0) | 68.2<br>(+0.0) | 71.5<br>(+0.0) | 69.1<br>(+0.0) | 78.9<br>(+0.0) | 73.5<br>(+0.0) | 72.1<br>(+0.0) | 69.6<br>(+0.0) | 75<br>(+0.0) | 62.6<br>(+0.0) |
|  | katG | S315T |  |  |  |  |  |  |  |  |  |  |
|  | inhA promoter | c -15 t |  |  |  |  |  |  |  |  |  |  |
| Clinical isolate | katG | S315T | 76.2<br>(+0.0) | 68.2<br>(+0.0) | 71.5<br>(+0.0) | 68.9<br>(±0.1) | 76.2<br>(+0.0) | 70.3<br>(±0.1) | 71.2<br>(+0.0) | 69.5<br>(±0.1) | 74.9<br>(+0.0) | 64.1<br>(+0.0) |
|  | eis promoter | g -10 a |  |  |  |  |  |  |  |  |  |  |
| Clinical isolate | katG | S315T | 76.4<br>(±0.1) | 68.3<br>(±0.1) | 71.6<br>(+0.0) | 69.2<br>(+0.1) | 76.3<br>(±0.1) | 70.4<br>(+0.0) | 71.3<br>(+0.0) | 69.7<br>(+0.0) | 75.1<br>(±0.1) | 64.2<br>(+0.0) |
|  | eis promoter | g -10 a |  |  |  |  |  |  |  |  |  |  |
| Clinical isolate | katG | S315T | 71.1<br>(+0.1) | 68.4<br>(+0.1) | 71.7<br>(+0.1) | 69.1<br>(+0.1) | 76.5<br>(+0.1) | 70.3<br>(+0.1) | 71.4<br>(+0.0) | 69.7<br>(+0.1) | 75.2<br>(+0.1) | 62.7<br>(+0.0) |
|  | inhA promoter | c -15 t |  |  |  |  |  |  |  |  |  |  |
|  | eis promoter | c -12 t |  |  |  |  |  |  |  |  |  |  |
| Clinical isolate | katG | S315T | 70.9<br>(+0.0) | 68.2<br>(+0.0) | 71.5<br>(+0.0) | 69<br>(+0.0) | 76.3<br>(+0.0) | 70.2<br>(±0.2) | 71.4<br>(+0.0) | 69.6<br>(+0.1) | 75<br>(+0.1) | 62.6<br>(+0.0) |
|  | inhA promoter | c -15 t |  |  |  |  |  |  |  |  |  |  |
|  | eis promoter | c -12 t |  |  |  |  |  |  |  |  |  |  |
| ** Capital letters > amino acid change and small letters > nucleotide change |  |  |  |  |  |  |  |  |  |  |  |  |

### Xpert MTB/XDR assay has a Limit of Detection (LoD) that is comparable to Xpert MTB/RIF for *M. tuberculosis* detection

The Xpert MTB/XDR assay includes a separate call out for *M. tuberculosis* detection, which is based on the identification of an *inhA* probe Tm in either WT or mutant windows. Positive detection of *M. tuberculosis* is required before the assay software will generate a resistance “DETECTED” or ‘NOT DETECTED” call. Thus, if the *inhA* probe does not result in a detectable Tm (WT or mutant), the result will be “MTB NOT DETECTED”, and no resistance result output will be available irrespective of whether Tm values are generated from the other targets in the assay. The Xpert MTB/XDR assay is designed to be run as a reflex test after initial testing has identified the presence of *M. tuberculosis* in the sample. Thus, our preference was to ensure that the Xpert MTB/XDR assay was at least as sensitive as the Xpert MTB/RIF assay in its *M. tuberculosis* detection function. The LoD of *M. tuberculosis* for the Xpert MTB/XDR assay was determined by performing a head-to-head comparison with the Xpert MTB/RIF assay, using the same *M. tuberculosis* H37Rv stock cultures. Six concentrations (200, 100, 80, 60, 20 and 10 CFU/mL) of *M. tuberculosis* strain H37Rv mc^2^6030 were spiked into pooled sputum samples confirmed to be *M. tuberculosis* negative by the Xpert MTB/RIF Ultra assay and tested in parallel by the Xpert MTB/RIF and the Xpert MTB/XDR assays in 20 replicates per concentration. The LoD calculated by Probit analysis was 71.9 CFU/mL (95% CI 58, 100) for the Xpert MTB/XDR assay and 86.9 CFU/mL 95% CI (72, 110) for the Xpert MTB/RIF assay (Figure 1). The LoD analyzed for each drug susceptibility call separately was 79.8 CFU/mL for INH, 95.5 CFU/mL for FLQ, 92.2 CFU/mL for AMK, 74.5 CFU/mL for KAN and 74.8 CFU/mL for CAP and 71.9 CFU/mL for ETH. Separate LoD studies were also performed at a different laboratory setting with two different lots of Xpert MTB/XDR cartridges and *M. bovi*s BCG stock instead of H37Rv to address the reproducibility of the initial LoD estimate. This study resulted in higher LoD estimates of 126-136 CFU/mL, when compared to the initial study using H37Rv stock, but was still comparable to the initially published Xpert MTB/RIF LoD (37). This minor difference in LoD estimates may be attributed to differences in cartridge lots and two different CFU stocks used to generate contrived samples. When we tested the LoD on NALC-NaOH concentrated sputum samples using spiked *M. bovis* BCG stock, the LoD was observed to be 86 CFU/mL, which is similar to the LoD obtained with direct sputum using H37Rv mc^2^6030 stock. Both the studies with the *M. bovis* BCG stock are described in detail in the supplementary results section.

**Figure 1:**
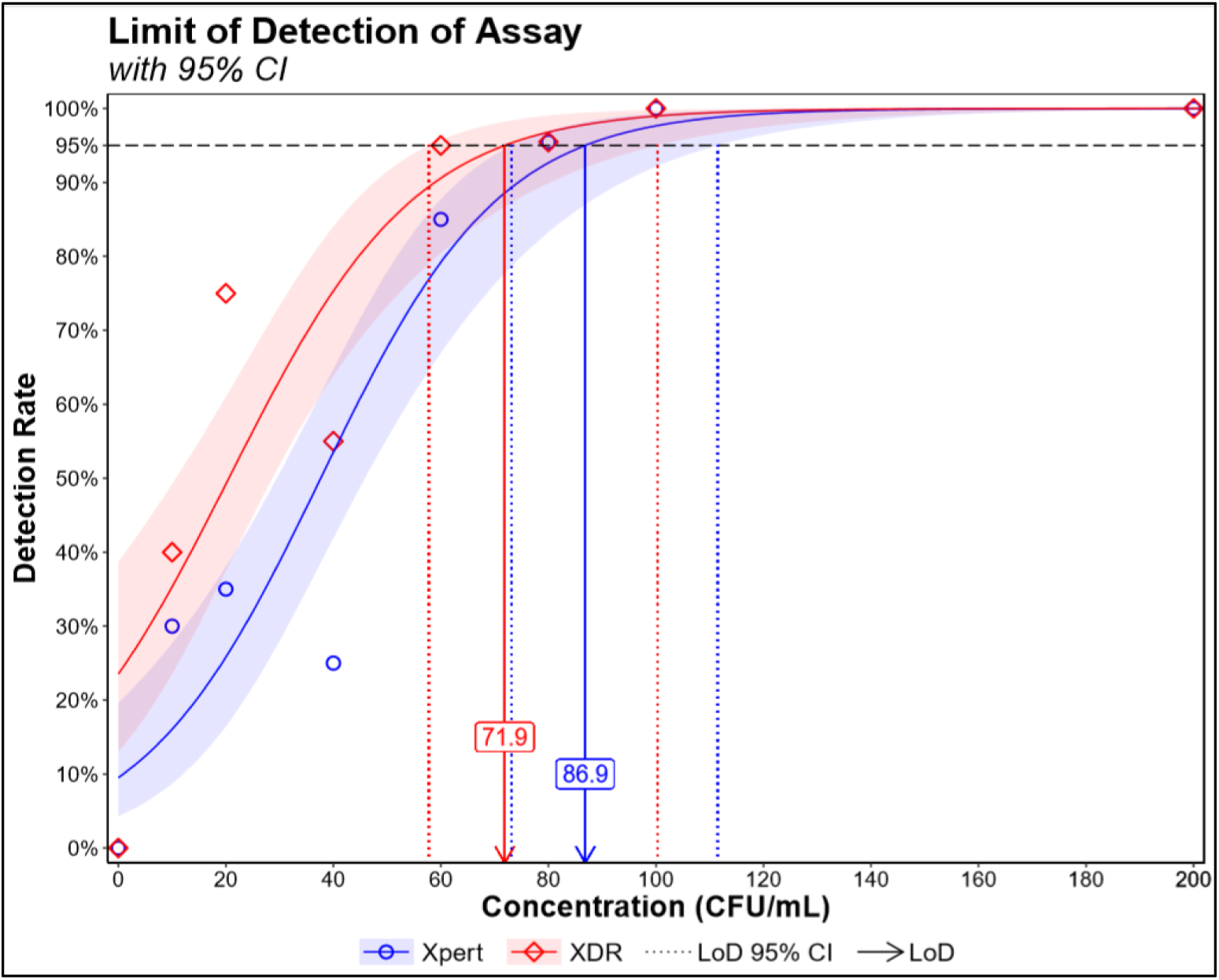
Limit of Detection of the Xpert MTB/XDR assay and the Xpert MTB/RIF assay performed side-by-side, with a minimum of 20 replicates for each cell concentration. Both assays were tested at 10, 20, 40, 60, 80, 100 and 200 CFU/mL and probit analysis was performed to calculate the LoD using R studio.

### Ability to detect a genetically diverse set of *M. tuberculosis* complex strains

To assess the capacity of the Xpert MTB/XDR assay to detect different species in the *M. tuberculosis* complex (MTBC), 9 *M. tuberculosis* complex species including *M. tuberculosis* strain H37Rv, *M. bovis, M. africanum, M. canetti*, and *M. microti* were tested by the Xpert MTB/XDR assay. These isolates were selected to be phylogenetically diverse with one representative strain across all major lineages of *M. tuberculosis* (26). All samples were detected as MTB positive (Table 3). *M. canetti* showed a *gyrB* Tm of 67.8°C which was a shift of −1.8°C from the mean WT Tm of 69.6°C as shown in Table 3 due to a polymorphism c/t in codon 533 in the *gyrB* gene present in some strains of *M. canetti* (27). However, this did not cause any false FLQ resistant calls since the *gyrB* Tm remained within the defined WT Tm window for *gyrB* probe. All of the other *M. tuberculosis* complex species tested generated WT Tm values identical to those of H37Rv and “Resistance NOT DETECTED” result output for all the drugs.

**Table 3.** Melting temperature values generated by the Xpert MTB/XDR assay tested for major *M. tuberculosis* complex lineages and species.

| RFLP | Lineage<br>(S.<br>gagneux) | Melt temperature (±SD) (°C) | | | | | | | | | |
| --- | --- | --- | --- | --- | --- | --- | --- | --- | --- | --- | --- |
|  |  | inhA | katG | fabG | ahp<br>C | gyrA1 | gyrA2 | gyrA3 | gyrB2 | rrs | eis |
| AH1 | Lineage 4 | 76.3<br>(±0.2) | 73.7<br>(±0.1) | 71.5<br>(+0.1) | 69.1<br>(+0.2) | 76.3<br>(+0.1) | 70.3<br>(+0.1) | 71.2 (+0.1) | 69.6<br>(+0.1) | 75 (+0.1) | 68.5<br>(+0.1) |
| HR36 | Lineage 5 | 76.4<br>(+0.2) | 73.7<br>(+0.1) | 71.5<br>(+0.1) | 69<br>(+0.1) | 76.3<br>(+0.1) | 70.3<br>(+0.1) | 71.2 (+0.1) | 69.6<br>(+0.2) | 75.0<br>(+0.1) | 68.6<br>(+0.1) |
| AR2 | Lineage 2 | 76.2<br>(+0.1) | 73.6<br>(+0.1) | 71.4<br>(+0.1) | 68.8<br>(+0.1) | 76.3<br>(+0.0) | 70.3<br>(+0.1) | 71.2 (+0.0) | 69.5<br>(+0.1) | 74.9<br>(+0.1) | 68.5<br>(+0.0) |
| GD139 | Lineage 3 | 76.3<br>(+0.2) | 73.7<br>(+0.1) | 71.5<br>(+0.1) | 70.3<br>(+0.2) | 76.4<br>(+0.1) | 70.3<br>(+0.1) | 71.2 (+0.1) | 69.6<br>(+0.2) | 75.0<br>(+0.2) | 68.6<br>(+0.1) |
| BCG | Lineage 1 | 76.2 | 73.6<br>(+0.1) | 71.5<br>(+0.0) | 68.6<br>(+0.0) | 76.3<br>(+0.0) | 70.3<br>(+0.1) | 71.15<br>(+0.1) | 69.5<br>(+0.0) | 74.9<br>(+0.0) | 68.5<br>(+0.0) |
| H37Rv | Lineage 4 | 76.3<br>(+0.0) | 73.7<br>(+0.0) | 71.4<br>(+0.1) | 69.3<br>(+0.0) | 76.1<br>(+0.1) | 70.3<br>(+0.1) | 71.1 (+0.1) | 69.6<br>(+0.0) | 75 (+0.0) | 68.5<br>(+0.1) |
| M. bovis | Animal<br>strain | 76.4<br>(+0.0) | 73.8<br>(+0.0) | 71.6<br>(+0.1) | 69.1<br>(+0.0) | 76.4<br>(+0.1) | 70.3<br>(+0.1) | 71.2 (+0.0) | 69.6<br>(+0.1) | 75.0<br>(+0.1) | 68.6<br>(+0.1) |
| M.<br>canetti | n/a | 76.4<br>(+0.1) | 73.8<br>(+0.0) | 71.6<br>(+0.0) | 69.3<br>(+0.1) | 76.4<br>(+0.0) | 70.3<br>(+0.1) | 71.3 (+0.1) | 67.8<br>(+0.0) | 75.1<br>(+0.1) | 68.6<br>(+0.0) |
| M.<br>microti | Animal<br>strain | 76.4<br>(+0.1) | 73.8<br>(+0.0) | 71.6<br>(+0.1) | 69.1<br>(+0.2) | 76.4<br>(+0.1) | 70.3<br>(+0.0) | 71.2 (+0.1) | 69.6<br>(+0.1) | 75.1<br>(+0.1) | 68.6<br>(+0.0) |

### Analytical specificity and cross reactivity

The specificity of the assay was assessed by testing 30 NTMs, 19 Gram-positive and Gram–negative bacteria, along with *Candida albicans* at 106 to 107 CFU/mL (Supplementary tables S1 and S2). All of the samples generated “MTB NOT DETECTED” results by the Xpert MTB/XDR assay, which specifically requires that the *inhA* probe generate a Tm in either the WT or MUT window to be called *M. tuberculosis*-positive. All of the NTM species tested, except *M. triviale*, generated *rrs* WT Tm values, which was expected because the target region of *rrs* gene is conserved among most Mycobacterium species (Supplementary table S1). The *rrs* WT Tm values were also obtained from *Citrobacter freundii, Corynebacterium xerosis, Enterobacter cloacae, Nocardia asteroids, Staphylococcus epidermidis, Streptococcus pyogenes* and *Candida albicans* indicating *rrs* primer/probe sequence overlap with the 16S ribosomal gene in these strains (Supplementary table S2). None of the strains was misidentified as *M. tuberculosis* positive due to the absence of any *inhA* promoter Tm. *M. gastri, M. gordonae*, and *M. xenopi* showed weak Tm peaks in the gyrA1 MutB Tm window and *M. interjectum* generated a gyrB2 WT Tm (Supplementary table S1). The rest of the targets did not cross-react in any of the NTMs. Very weak *gyrA* probe cross-reactivity was observed with a few additional bacterial species (Supplementary table S2).

Since weak *gyrA* mutant Tm peaks were observed for some of the NTM species, we performed spiking experiments with BCG and NTM mixtures to simulate the clinical scenario of a tuberculosis patient who is also infected with an NTM. Studies were then undertaken to test whether this type of dual infection could generate a false-positive FLQ resistance call. High concentrations (106 CFU/mL) of 12 clinically relevant NTM species were mixed with a low concentration of *M. bovis BCG* (408 CFU/mL, i.e. approximately 3X LoD) and tested with the Xpert MTB/XDR assay (Supplementary table S3). None of the NTMs tested in these mixtures generated a false FLQ resistance calls. However, we did observe that one strain of *M. marinum* (ATCC 0927) interfered with the *gyrA* signal produced by the *M. tuberculosis* target resulting in the suppression of the Tm generated by at least one of the *gyrA* probe resulting in a “FLQ Resistance INDETERMINATE” call. At 106 CFU/mL all the 4 replicates tested, generated Indeterminate calls for FLQ and at 105 CFU/ml, 2 of 4 replicates resulted in indeterminate calls. This suppression only occurred when samples were spiked with 105 *M. marinum* CFU/mL or above. With *M. marinum* ATCC 0927 spiked at 104 CFU/mL, no interference was observed and the correct “FLQ Resistance NOT DETECTED” call was observed. To the best of our knowledge, there have been no reports of pulmonary infections caused by co-infection with *M. tuberculosis* and *M. marinum* and thus this interference may not be clinically relevant, at least for pulmonary TB cases.

### Detection of hetero-resistance

It is estimated that about 20% to 45% of XDR cases contain a mixed population of susceptible and resistant strains, i.e. are hetero-resistant (28-31). We have shown previously that SMB assays can efficiently detect WT and mutant DNA mixtures by generating double Tm peaks corresponding to WT and mutant DNA sequences (10). To assess the performance of the Xpert MTB/XDR assay to detect mutations in hetero-resistant samples, *E. coli* cells transfected with plasmids containing WT or mutant XDR target sequences were used. A series of cell mixtures containing 0%, 10%, 15%, 20%, 25%, 50%, 60%, 75%, 90% and 100% of mutant was tested against a background of cells with WT sequences in replicates of three. The total amount of cells tested in each mixture was fixed at 10,000 cells/mL. We used this approach to test mixtures of WT cells and cells containing the mutations: c(−15)t in the *inhA* promotor, S513T in the *katG* gene, g609a in *fabG1*, c(−39)t in *oxyR-ahpC* region, D94G in the *gyrA*, E540D in *gyrB*, a1401g in *rrs*, and c(−14)t in the *eis* promoter. Resistance was detected when the mutant Tm could be detected in presence of the WT background and “Resistance NOT DETECTED” calls were made when the mutant Tm was undetectable and only WT Tm was detected (Figure 2). For detection of *fabG1, katG* or *inhA* promoter mutations, our results showed that INH-R was detected in mixtures containing as little as 20% mutant cells in 80% of WT cells. However, cells containing an *ahpC* mutant could not be detected unless they were present in at least 75% of the mixture. FLQ-R was detected in mixtures that contained as little as 25% of the D94G mutation; however, mixtures containing the *gyrB* mutation were only detected in mixtures that contained 60% or more of the mutant sequence. SLID resistance was detected in mixtures that contained as little as 60% of the *rrs* or the *eis* promoter mutations. Below these levels, resistance was not detected, since the mutant Tms could not be identified against the WT Tm background as shown in figure 2.

**Figure 2:**
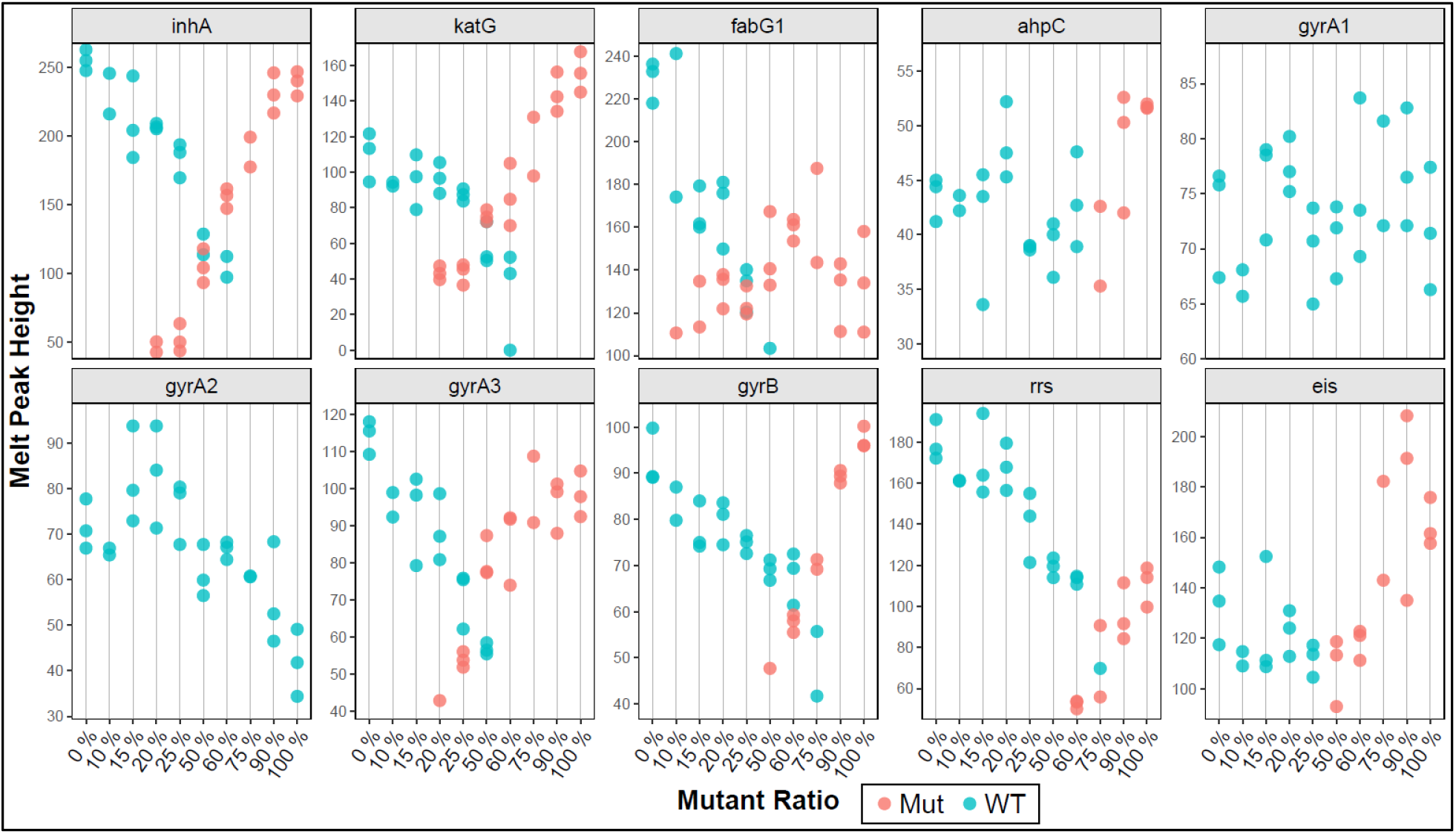
Melt peak heights of each target in mixtures containing different ratios of cells containing WT and mutant plasmids respectively, where blue dots indicate a susceptible call and red dots indicate mutant calls based on their Tm and melt peak height. The melt peak height is determined by highest distance between the peak of the first derivative melt curve and baseline. The presence of blue and red dots for any concentration designates detection of both a WT and a mutant Tm. In such cases the results obtained was “RESISTANCE DETECTED” for the corresponding drug. The QRDR mutation D94G generates a mutant Tm only with the gyrA3 probe. Each mixture was tested in triplicate, except the 10% and 75% mixtures, which had two valid replicates.

We also evaluated the ability of the assay to accurately detect the three low FLQ resistance-associated mutations, A90V, S91P and D94A, when present as a mixture with WT sequence. We tested three replicates from each mixture, containing 0%, 10%, 20%, 30%, 40%, 50% and 100% mutant cells in a background of WT cells in a total mixture of 5000 cells. We found that the assay was not able to detect low FLQ resistance for all the A90V and S91P mixtures we tested and generated either a “FLQ Resistance DETECTED” call or “FLQ Resistance NOT DETECTED” call. In the former case, the presence of Tm values in both WT and MUT windows produced a Tm pattern that was consistent with “FLQ Resistance DETECTED” call, and in the latter case, there was only WT Tm present. A “FLQ Resistance DETECTED” call was made for S91P/WT mixtures down to as little as 20% S91P. Below that concentration, S91P/WT mixtures produced a “FLQ Resistance NOT DETECTED” call, since no mutant Tm could be detected. A “FLQ Resistance Detected” call was made for A90V/WT mixtures down to as little as 20% - 50% A90V and mixtures with less than 20% A90V produced a “FLQ Resistance NOT DETECTED” call. In contrast, with the D94A mutation, the assay was able to correctly detect low-level FLQ resistance in hetero-resistant samples containing at least 50% D94A mutant cells mixed with 50% WT cells in all three replicates tested, due to the correct D94A specific Tm signature (MutA-MutA-WT) being present. Two of three replicates containing 40% D94A produced a “Low FLQ Resistance DETECTED” call and one replicate produced “FLQ Resistance DETECTED” call. At D94A/WT mixtures below 40% D94A, the assay was unable to detect the presence of FLQ specific mutations (Data not Shown). Thus, the assay demonstrates a substantial loss in the ability to distinguish low-FLQ resistance conferring A90V and S91P mutations from other QRDR mutations, when these two mutations are present along with WT sequence, but its overall ability to identify FLQ resistance is not affected.

### Performance on sputum samples and clinical isolates

A limited clinical study was performed at two different testing sites with a total of 100 *M. tuberculosis* positive frozen sputum and 214 clinical isolates from de-identified patients with XDR TB. The sensitivity and the specificity of the assay for detecting resistance to INH, ETH, FLQ and SLIDs on this sample set were estimated by individually comparing to the two different reference standards: P-DST and DNA sequencing. The capacity of the assay to accurately detect the mutations in the target genes was also estimated. The results from 105 of 107 clinical isolates and all the 50 sputum samples were available from site 1 and the results from 106 out of 107 isolates and 49 of the 50 sputum samples were available from site 2, which allowed us to include 310 of the 314 samples for the final analysis. Any “Indeterminate” result for any drug targets and samples with missing or ambiguous P-DST and/or sequencing results were excluded from the analysis. Excluding such samples, P-DST results were available for 309 samples for INH, 306 samples for KAN, 305 samples for each of FLQ and CAP, 303 samples for AMK and 265 samples for ETH. Similarly, sequencing results were available for all the 310 samples for INH and ETH specific targets, 309 samples for FLQ specific targets, 306, 307 and 308 samples for AMK, CAP and KAN specific targets respectively. Compared to P-DST, the assay showed a sensitivity and specificity respectively of 98.3% and 95% for INH resistance, 91.4% and 98.5% for FLQ resistance, 91% and 99% for AMK resistance, 98.1% and 97% for KAN resistance, 70% and 99.7% for CAP resistance and 65.4% and 97.3% for ETH resistance (Table 4). Compared to sequencing, the assay showed sensitivity of 99.7%, 97.5%, 100%, 96.5%, 94.1% and 88.5% for detection of INH, FLQ, AMK, KAN, CAP and ETH resistance respectively, with a specificity of 100% for all the drugs except for ETH with a specificity of 97.3% (Table 4). In this sample set, mutations present in the key target genes were as follows; g(−17)t, c(−15)t, t(−8)a and t(−8)c in *inhA* promoter region, S315T, S315N and S315G in the *katG* gene, L203L in the *fabG1* gene, G88C, D89N, A90V, D94A, D94G, D94Y in the *gyrA* gene, “a1401g” in the *rrs* gene and g(−37)t, c(−14)t, c(−12)t, g(−10)a and c(−8)a in the *eis* promoter region. All the *gyrB* and *ahpC* mutations present in this sample set were associated with mutations in the *gyrA* gene and *inhA* promoter or *katG* genes, respectively. The assay detected all the mutations present in the *inhA* promoter region and the *katG* gene, as well as all the low FLQ resistant A90V and D94A mutations and differentiated them from high FLQ resistance caused by other *gyrA* QRDR mutations. The assay also correctly detected SLID cross-resistance and individual resistance to KAN by correctly identifying the mutations in the *rrs* and *eis* promoter genes respectively. As shown in Figure 3, the assay was able to clearly and independently cluster the WT and mutant Tm values for all the targets resulting in unequivocal identification of these mutations with a high degree of accuracy. A very few “Indeterminate” results were obtained for FLQ (0.3%), AMK (1.3%), KAN (0.6%) and CAP (0.9%) due to missing Tm values from one or more of the key analytes related to detection of resistance to these drugs. We observed that in all these “Indeterminate” calls, the respective Tm peaks were present, but they were not high enough to cross their pre-defined Tm peak height threshold, and thus the Tm values were not calculated by the assay algorithm, which indicates possible sub-LoD concentrations of the targets. In at least one case of CAP Indeterminate results, the missing Tm from *rrs* probe could be attributed to unexpected optical aberrations in the instrument module, which prevented determination of the melt peak.

**Table 4.** Xpert MTB/XDR assay’s concordance with P-DST and sequencing on drug resistance detection on clinical isolates and sputum samples

|  | **P-DST** |  |  |  |  |  |  |  |  |
| --- | --- | --- | --- | --- | --- | --- | --- | --- | --- |
| **Drugs** | **N** | **TP** | **FN** | **TN** | **FP** | **Sensitivity (%)** | **95%CI** | **Specificity (%)** | **95%CI** |
| **INH** | 309 | 284 | 5 | 19 | 1 | 98.3 | 95.8-99.3 | 95.0 | 73.1-99.7 |
| **FLQ** | 305 | 32 | 3 | 266 | 4 | 91.4 | 78.9-98.9 | 98.5 | 95.9-99.5 |
| **AMK** | 303 | 20 | 2 | 278 | 3 | 91.0 | 69.4-98.4 | 98.9 | 96.6-99.7 |
| **KAN** | 306 | 101 | 2 | 197 | 6 | 98.1 | 92.5-99.7 | 97.4 | 93.4-98.8 |
| **CAP** | 305 | 14 | 6 | 284 | 1 | 70.0 | 45.6-87.2 | 99.7 | 97.7-99.9 |
| **ETH** | 265 | 102 | 54 | 106 | 3 | 65.4 | 57.3-72.7 | 97.3 | 91.6-99.3 |
|  | **Sequencing** |  |  |  |  |  |  |  |  |
| **Drugs** | **N** | **TP** | **FN** | **TN** | **FP** | **Sensitivity (%)** | **95%CI** | **Specificity (%)** | **95%CI** |
| **INH** | 310 | 286 | 1 | 23 | 0 | 99.6 | 97.8-99.9 | 100 | 82.2-100 |
| **FLQ** | 309 | 39 | 1 | 269 | 0 | 97.5 | 85.3-99.8 | 100 | 98.2-100 |
| **AMK** | 306 | 24 | 0 | 282 | 0 | 100 | 82.8-100 | 100 | 98.3-100 |
| **KAN** | 308 | 109 | 4 | 195 | 0 | 96.4 | 90.6-98.9 | 100 | 97.6-100 |
| **CAP** | 307 | 16 | 1 | 290 | 0 | 94.1 | 69.2-99.6 | 100 | 98.4-100 |
| **ETH** | 310 | 108 | 14 | 183 | 5 | 88.5 | 81.2-93.4 | 97.3 | 93.6-99.0 |

N= Number of samples; TP: True Positive; FN: False Negative; TN: True Negative; FP: False Positive

**Figure 3:**
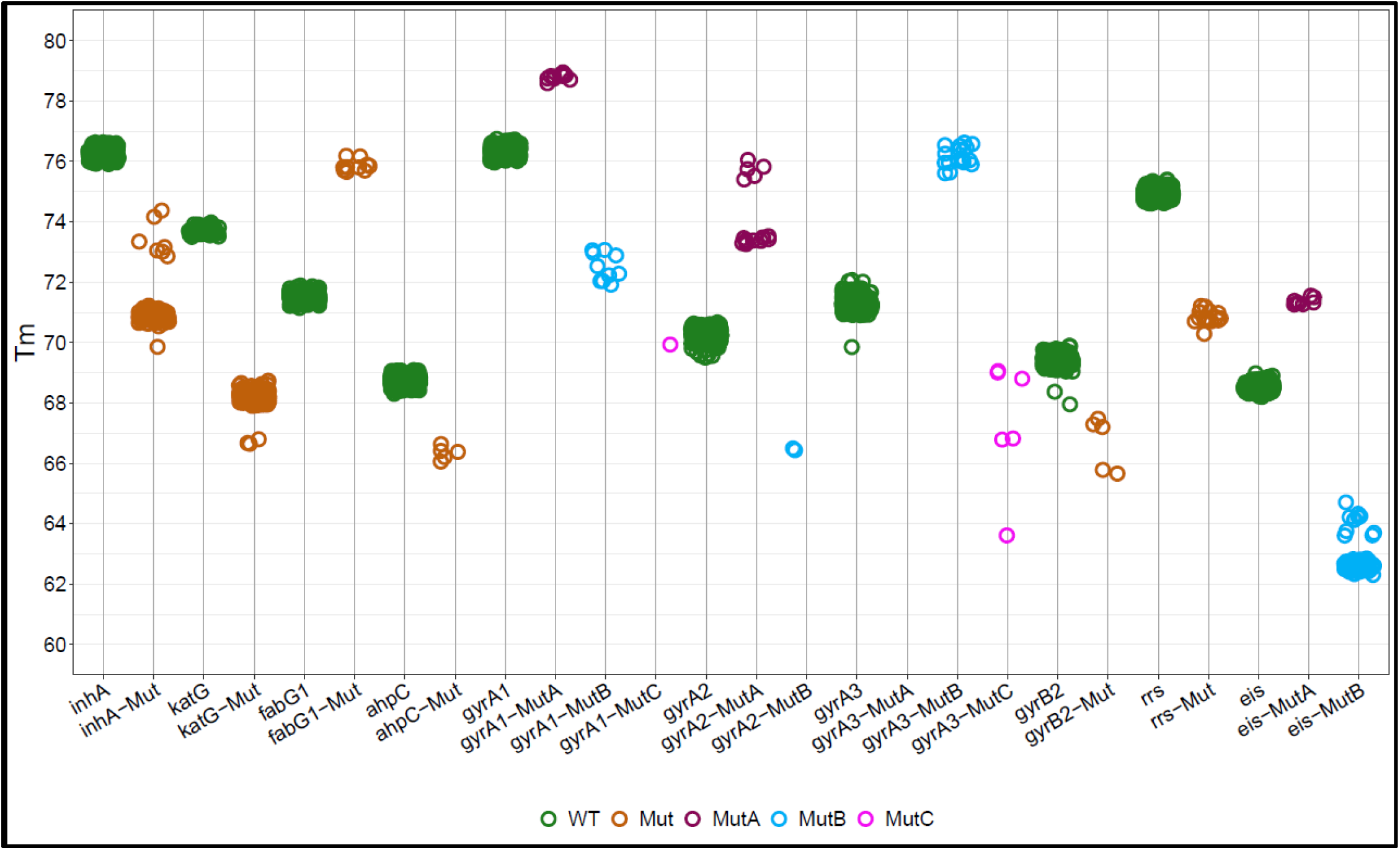
Scatterplot showing clustering of WT and mutant Tm values from Clinical study with 100 *M. tuberculosis* positive frozen sputum and 214 clinical isolates for all Xpert MTB/XDR targets using ggplot on R studio. To prevent over plotting, a degree of jitter was introduced. All green dots are WT Tm while brown, purple, blue and pink are mutant Tm values.

## Materials and Methods

### Description of the assay

The Xpert MTB/XDR assay is a 9-plex assay consisting of 10 sloppy molecular beacon (SMB) probes (32) that target 8 different *M. tuberculosis* genes detecting resistance to INH, ETH, FLQ and SLID. To detect INH resistance, four probes target the *inhA* promoter (nucleotides −1 to −32), the *katG* (codons 311-319) and *fabG1* (codons 199-210) genes, and the *oxyR*-*ahpC* (*ahpC*) intergenic region (nucleotides −5 to −50). Identification of *inhA* promoter mutations in a specific optical channel additionally enables detection of ETH resistance and allows differentiation of low-level INH resistance, as both resistance characteristics are encoded by mutations in the *inhA* promoter (33-35). To detect FLQ resistance, three overlapping probes target the *gyrA* QRDR (codons 87-95) and one probe targets the *gyrB* QRDR (codons 531-544). The three *gyrA* probes used in the assay have 8 defined mutant windows, which enables specific identification of the QRDR mutations A90V, S91P and D94A associated with low-level FLQ resistance and differentiates them from other QRDR mutations associated with higher-level FLQ resistance, as described in detail in the results section. To detect SLID resistance, namely AMK, KAN and CAP, one probe targets the *rrs* gene (nucleotides 1396-1417) and a second probe targets the *eis* promoter region (nucleotides −6 to −42). Cross-resistance between AMK, KAN and CAP is well documented (15-17) and is captured by *rrs* probe. The *eis* probe can differentiate between c(−14)t which confers cross-resistance to KAN and AMK and other mutations in the *eis* promoter region which only confer resistance to KAN, since the probe has been specifically designed to generate a higher Tm shift for the c(−14)t mutation and a lower Tm shift for all the other mutations from the WT Tm. To serve as an internal sample processing and PCR control, an additional SPC probe targets a *Bacillus globigii* gene using the same fluorophore as the *ahpC* probe. This enabled us to accommodate the 11 probes in the assay within the 10 optical channels). Dehydrated *B. globigii* spore beads are included in the assay cartridge. The SPC probe is a Taqman probe, is only detectable during PCR amplification, and does not generate a melting curve and thus do not interfere with the melt signals from the *ahpC* SMB probe. Each of the 10 probes in the assay has one defined wild type (WT) Tm range (defined as a WT Tm window) and one or several Tm ranges that define the presence of mutants (mutant Tm windows). WT or mutant sequences are identified by the WT and mutant Tm values respectively for each target, which results in the “Resistance NOT DETECTED” or “Resistance DETECTED” calls respectively, for the related drugs. The PCR assay consists of two phases. The first phase is a conventional symmetric PCR, followed by a second nested asymmetric PCR phase, except for the SPC assay, which is symmetric in both phases. The nested asymmetric assay enables preferential amplification of the target strands to which the SMB probes bind with high efficiency even in this 9-plex assay system. Specific in-cartridge microfluidics allow the products of the first PCR to be added to the second PCR after 31 cycles are complete. The second PCR consists of 40 cycles followed by the melt curve stage. No third stage of linear PCR is used in this assay unlike the earlier XDR cartridge, which reduces the time to result from 120 to under 90 minutes.

### Cartridge configuration, assay composition and testing procedure

The Xpert MTB/XDR cartridge is a modified version of the prototype assay cartridge described previously (36). It consists of a multi-position fluidic valve, a bacterial capture chamber and 11 chambers containing buffers and reagents for sample processing and PCR, plus an integrated 50uL PCR tube. Two sets of two lyophilized beads each are used to amplify resistance-conferring regions of *inhA* promoter, *katG, fabG1, ahpC, gyrA, gyrB, rrs* and *eis* promoter as well as an internal control sequence of *B. globigii*. The first bead set is used to perform 9-plex PCR, followed by a full nested or hemi-nested PCR of the first set of amplicons using the second bead set. The second bead set contains SMBs for 10 gene targets and a Taqman probe for the internal control. The SMBs for the *ahpC* target and internal control share the same channel and are designed not to interfere with other’s detection.

To perform a test, each sample (spiked sputum, clinical sputum samples, cultured *M. tuberculosis* or *M. bovis* BCG CFU) was first mixed at a 2:1 ratio with an NaOH and isopropanol containing sample reagent (SR) as described previously(37); the sample was then added to the sample loading chamber of the cartridge (for CE-IVD use, sputum is the only recommended sample type currently for diagnostic purposes). The loaded cartridge was placed into a GeneXpert instrument running software developed for the Xpert MTB/XDR assay (Cepheid, Sunnyvale, CA). The assay was then started and automated processing of the sample for DNA isolation followed by the two-phase PCR assay and melt analysis was performed. The microfluidics were similar to that previously described (36). Briefly, the internal control *B. globigii* spores were mixed with the sample, and the sample was then filtered through a bacterial capture filter within the cartridge. The captured bacteria and *B. globigii* spores were then washed multiple times with a wash buffer and DNA released from captured bacteria by highly efficient lysis through sonication. The lysate was then used to re-suspend the first set of PCR reagent beads. The PCR mix was aspirated into the PCR tube to initiate the first round of amplification. After this first PCR, a specific amount of the amplified sample was moved out of the tube and the PCR tube was then filled with a second set of PCR reagent beads that had been re-suspended in buffer and mixed with the remaining amplicon in the tube. A second PCR was then performed followed by a post PCR melt analysis, to generate first derivative Tm curves. The Tm values were identified by the automated GeneXpert Tm calling software (Cepheid, Sunnyvale, CA) and classified as Tm values that identified WT or mutant amplicon sequence based on pre-defined Tm parameters (Tm windows). These Tm values were then used to determine the presence or absence of resistance to the target drugs.

### Preparation of *M. tuberculosis* and *M. bovis* BCG culture stocks and determination of CFU

*M. tuberculosis* culture stock preparation and CFU counts were determined as described previously (36). An attenuated strain of *M. tuberculosis* H37Rv (mc^2^6030) or *M. bovis* BCG was cultured by inoculating 1:100 in 20 mL of Difco Middlebrook 7H9 media (BD Biosciences, California USA) supplemented with 10% BBL Middlebrook OADC Enrichment (BD Biosciences, California USA), 0.05% Tween 80 (Sigma Aldrich, St Louis, MO) and 24 μg/mL Calcium Pantothenate (Sigma Aldrich, St Louis, MO) for *M. tuberculosis* H37Rv (mc^2^6030). The strains were grown to the optical density 600nm of 0.6-0.8, sub-cultured twice before performing dilutions for CFU determinations and storage. The cultures were quantified by plating 10-5, 10-6 and 10-7 dilutions in triplicate on 7H10 plates supplemented with 10% OADC and 24 μg/mL Calcium Pantothenate fo*r M. tuberculosis* H37Rv (mc^2^6030). The cultures were divided in 500 uL and stored at −80°C until use. Colony counts was performed 3 weeks after plating, once the colonies became visible.

### Dilutions and spiking in sputum for determining Limit of Detection

To dilute and spike *M. tuberculosis* or *M. bovis* BCG in sputum for analytical studies, a frozen aliquot was processed as previously described (36). Briefly, the frozen aliquot was thawed at room temperature and re-suspended in 7H9 media up to 1 mL. The aliquot was then vortexed for 30 seconds and allowed to rest on ice for 6 minutes. The aliquot was then sonicated for 30 seconds using Branson CPX1800 Ultrasonic water-bath (Fisher Scientific, Waltham, MA, USA), followed by standing on ice for 30 seconds. This step was repeated twice more followed by resting for 6 minutes on ice. The sonicated stock was then used to prepare the required dilutions in 4 mL 7H9 media up to 1000 CFU/mL. Controlled mixing steps were performed during serial dilutions by aspirating and dispensing via pipette. The sonicated aliquot was stored at 4°C for no more than 7 days. If the sonicated aliquot was used on subsequent days, it was sonicated once for 30 seconds when re-used after 24 hrs. of first storage and 2 minutes of vortexing for any subsequent dilution experiments. The 1000 CFU/mL dilution was spiked in sputum to obtain the final test concentration. SR was added at 2:1 ratio to allow sufficient volume to distribute in 2 mL in each cartridge. To determine LoD, we tested 10, 20, 40, 60, 80, 100, 200 CFU/mL, 20 replicates each. For detection of LoD in concentrated sputum, unprocessed sputum samples were first spiked with the target level of *M. bovis* BCG CFU and each spiked sputum specimen was processed to obtain concentrated sediment using the BD BBLTM MycoprepTM mycobacterial system digestion/decontamination kits (Becton Dickinson, Franklin Lakes, NJ, USA) following manufacturer’s instructions. LoD was calculated by probit analysis performed on R studio.

### Detection of *gyrA* QRDR mutations

We have created a repository of plasmids by cloning approximately 100 bp fragment of *gyrA* gene in vector p.MV306H with the help of gene synthesis and mutagenesis services of Genscript Biotech Inc. (Piscataway, NJ, USA). The cloned fragments contained the individual *gyrA* QRDR single point mutations that are frequently associated with FLQ resistance (18-20) in each plasmid. Lyophilized recombinant plasmids were re-suspended in water and quantified by NanoDrop-1000. To determine Tm of *gyrA* probes, each plasmid was tested multiple times at 100 pg/rxn. Mean Tm and standard deviations were calculated using Microsoft Excel 365.

### Preparation of mixed cells to test for detection of heteroresistance

Quantified preparations of hardened *E. coli* cells (Maine Molecular Quality Controls Inc., Saco, Maine, USA) that had been transfected with plasmids containing WT or mutant target sequences including c(−15)t in the *inhA* promotor, S513T in the *katG* gene, L203L in *fabG1*, c(−39)t in *oxyR-ahpC* region, D94G in the *gyrA*, E540D in *gyrB*, a1401g in *rrs*, and c(−14)t in the *eis* promoter were used. A series of cell mixtures containing 0%, 10%, 15%, 20%, 25%, 50%, 60%, 75%, 90% and 100% of mutant was tested against a background of cells with WT sequences. The total amount of cells tested in each mixture was 10,000 cells/mL containing enough volume to test three replicates. Similarly, mixtures of low level FLQ resistance conferring mutations A90V, D94A and S91P in *gyrA* gene were prepared by mixing a series of cells containing 0%, 10%, 20%, 30%, 40%, 50% and 100% of mutant cells with cells with WT *gyrA* gene sequence at final cell counts 5000 cells/mL, replicates of three. SR reagent was added to each mixture at a 2:1 ratio and incubated for 15 minutes. 2 mL of the inactivated mixture was added to the cartridge. The cartridge placed into the GeneXpert instrument module and the assay performed by selecting a version of an automated assay protocol. Mean Tms were calculated and standard deviations were calculated in Microsoft Excel.

### Mutant panel challenge

DNA was extracted from a panel of 14 clinical isolates with canonical mutations in the target genes and promoter regions known to be associated with clinical INH, FLQ, ETH and SLID resistance, by a boiling preparation using InstaGene Matrix (Bio-rad, Hercules, CA USA) or using Phenol Chloroform method, each described previously (32, 38). All the mutations were confirmed by Sanger sequencing of each of the target genes and promoter regions. All isolates were quantified using the Qubit dsDNA HS Assay kit (ThermoFisher Scientific, Waltham, MA USA). All isolates were tested at a concentration that was approximately 3X of a predetermined BCG DNA LoD (in terms of genome equivalents), or higher when we observed indeterminate resistance calls resulting from absence of Tms due to poor quality of isolated DNA. The diluent solution (Tris EDTA Tween buffer) was used as Negative control, and BCG DNA at 3x the LoD was used as a positive control. Each isolate was tested in replicates of three. Mean Tms and standard deviations were calculated in Microsoft Excel. The isolates were pre-loaded in the XDR cartridge and the cartridge placed into the GeneXpert instrument module and the assay performed by selecting a version of an automated assay protocol that was slightly modified to permit testing of DNA rather than *M. tuberculosis* CFU.

### Analytical specificity and cross reactivity

Thirty species of Non-tuberculosis Mycobacterium (NTM), either purchased from the American Type Culture Collection (ATCC) or kindly provided by the National Jewish Health (Denver, CO, USA), were cultured and quantified similar as described for *M. tuberculosis* in 7H9 media supplemented with 10% Middlebrook OADC Growth supplement, and 0.05% Tween 80. We were able to cultivate and quantify all 30 NTM species except for *M. genevanse*, for which we could only record the optical density. Gram-negative and Gram-positive bacteria obtained from the University Hospital Microbiology Lab (University Hospital, Newark NJ, USA) were cultivated on blood agar plates. DNA was isolated by boiling preparation using an Instagene Matrix (32). The isolated DNA was measured by the Qubit dsDNA HS Assay Kit. Both NTM and Gram-negative and Gram-positive bacteria were tested at final concentration equivalent to 106 to 107 CFU/mL. Dilution buffer was used as the negative control and BCG (cells and DNA) at 3x LoD was used as the positive control.

Twelve clinically relevant NTMs were identified to test in a mixture with *M. bovis BCG* to identify possible interference for *M. tuberculosis* detection by high concentrations of NTMs. The NTMs were diluted to a final test concentration 106 CFU/mL and mixed with *M. bovis BCG* at a final test 3x LoD of the stock with enough volume for four replicates. Dilution buffer was used as the negative control and *M. bovis* BCG, at 3x LoD concentration, was used as the positive control. To all test samples, SR reagent was added and incubated for 15 mins. 2mL sample was aliquoted to each cartridge and loaded into the GeneXpert instrument.

### Clinical study protocol

A small clinical study was performed with well-characterized and archived frozen sputum and culture isolates from de-identified XDR TB patients provided by the Foundation of Innovative New Diagnostics (FIND) (39). The samples were obtained from Georgia, Moldova, Peru and Vietnam representing three different continents and were chosen to represent all the common clinically relevant mutations present in the target genes considering the global estimate of prevalence. The study was performed at two different sites at New Jersey Medical School, Rutgers University, Newark, NJ, USA and San Raffaele Hospital, Milan, Italy. The study consisted of 214 clinical isolates and 100 sputum samples from XDR TB patients, which were equally distributed between both the sites (107 clinical isolates and 50 sputum at each site). P-DST results were available for INH, ETH, at least one or more of the FLQs (Ofloxacin, Levofloxacin and Moxifloxacin) and the SLIDs (AMK, CAP and KAN). Sanger sequencing results were available for *katG, inhA* promoter, *fabG1, oxyR-ahpC* intergenic region, *gyrA, gyrB, rrs* and *eis* promoter i.e. all the target genes in the assay along with several other non-target genes associated with resistance to other first line drugs (including *pncA, ethA, ethA* upstream, *rpsl, tlyA, ndh* etc.). Each sputum sample consisted of 0.5 mL duplicate aliquots, which were thawed at room temperature and pooled, and vortexed thoroughly to ensure homogenization at each site before processing. The pooled 1 mL sputum was transferred to a new tube and 2 volumes of sample reagent (SR) was added to it, mixed and incubated for 15 minutes before performing the test. The frozen clinical isolates contained approximately 400-500µl of cell suspension at approximately 106 CFU/mL concentration. The isolates were thawed and sterile Tris pH 8.0 or PBS was added until the total volume of cell suspension was equivalent to 1 ml, vortexed well for 30 second and followed by standing for 5 minutes. Two volumes of Sample Reagent (SR) to the 1.0mL suspension, mixed and incubated for 15 minutes before performing the test. All the operators at each site as well as personnel performing the result interpretation and data analysis were completely blinded to the sequencing and the P-DST results. The aim of this study was to determine the diagnostic performance (sensitivity and specificity) of the Xpert MTB/XDR assay for INH, ETH, FLQ and SLID resistance detection compared to the individual reference standards P-DST (phenotypic reference standard) and sequencing (molecular reference standard). Analysis was performed by combining both the clinical isolates and sputum for a composite analysis for both the sample types. Additionally, “Indeterminate” rates for each drug type and “Non determinate” run (run aborts due to errors) rates of the assay for each sample type at each site were also calculated. The “Non determinate” samples were repeated only for the isolates, since a second aliquot was available. No repeat runs were performed for “Non determinate” sputum since the entire sample was used for the first run.

Statistical Analysis: LoD was calculated using the percentages of the replicates resulting in successful TB detection and drug susceptibility calls at each input CFU concentration in sputum for both Xpert MTB/XDR and Xpert MTB/RIF assays using probit analysis on R studio version 1.2.5019, “Elderflower” (RStudio Team (2020). RStudio: Integrated Development for R. RStudio, PBC, Boston, MA URL http://www.rstudio.com/). Binary probit regression results were fitted through the tested concentrations, and lower and upper 95% confidence intervals (95% CIs) were generated for the curve. The 95% CI for the minimum input concentration was determined by where the 95% probability level crossed the upper and lower 95% CIs, which indicated the LoD. Mean Tm and standard deviations were calculated on Microsoft Excel.

## Discussion

We describe here the development of an *in vitro* diagnostic version of the prototype cartridge for INH, FLQ and SLID resistance detection (10) with better coverage for INH resistance, the capacity to detect low versus high level resistance for INH and FLQ, to identify individual versus cross-resistance to SLID, as well as with better analytical sensitivity and a reduced time to result. The SMB probe design and chemistry of the new Xpert MTB/XDR assay is similar to the prototype cartridge, with additional probes added to detect new targets indicative of INH and ETH resistance, an additional probe and modified probe designs for the *gyrA* gene to differentiate and identify low vs high-level FLQ resistance, and a modified *eis* probe design to identify KAN resistance only, versus KAN/AMK cross resistance. An additional *gyrB* probe in the prototype version, which targeted the codon 500, was removed from this assay to accommodate the new probes on account of very low frequency of any high confidence mutations present in the codon 500 of *gyrB* (24). This modified assay version eliminates the three-stage PCR amplification used in the prototype cartridge and closely approximates the PCR cycling strategy used in the Xpert MTB/RIF Ultra assay, where the first stage of symmetric PCR is followed by a second stage of asymmetric PCR preferentially amplifying the target strands, followed by a melt stage. We have also utilized the strategy of including a Taqman probe (SPC) and an SMB probe (*ahpC*) with the same fluorophore to emit signal in a single channel, which allowed us to develop a 10 color, 11-probe assay. The *ahpC* probe was designed to have a probe-target hybrid Tm close to the annealing temperature of the assay, while the Taqman probe was designed to have a probe-target hybrid Tm at least 50C above the annealing temperature. This allowed us to generate good real time signals preferentially from the SPC Taqman probe during the amplification stage and obtain clear melt curves from the *ahpC* SMB probe during the melt stage, without any interference to melt signal from the Taqman probe.

Designing probes to distinguish high and low-level FLQ resistance was especially challenging, since we had to ensure that the Ser/Thr polymorphism at codon 95 was not recognized as a mutation, while all of the three low-level FLQ resistance-inducing mutations were individually identified and differentiated from all of the other QRDR mutations. Several iterative probe designs were tested based on the probes present in our prototype cartridge and a combination of three overlapping probes were chosen to generate a series of Tm signatures, which not only individually identified the three different mutations and differentiated them from other QRDR mutations, but also generated the same WT Tm values for the codon 95 polymorphism. We successfully used this Tm signature principle to identify these mutations by placing the mutant Tm values in carefully chosen, specific WT and mutant Tm windows for each probe, which underscores the previously described capacity for SMB probe tiling to accurately identify DNA sequences (40). The Xpert MTB/XDR assay targeted at least two regions in the *M. tuberculosis* genome which contained mutations spread over relatively long stretches, which would be very difficult for a single probe to query. These regions were the *oxyR-ahpC* intergenic region where mutations were spread over 46 bp and the *eis* promoter region where mutations were spread over 37 bp. We used poly dT and poly dA to link two different probes for the *ahpC* target region to generate a 49 bp long probe, and we used special proprietary Cepheid linkers to combine together two of the *eis* probes from the prototype cartridge assay to create a 50 bp long probe (including the linker sequence). We introduced mismatches in the probes to enhance the delta Tm between the WT and mutant sequences, to ensure that clearly separated Tm values were generated between mutant and WT sequences. These and other probe design principles were used to ensure that there was a ≥20C separation between WT and mutant Tm values for most clinically relevant mutations.

As a reflex INH and second line resistance detection assay to the Xpert MTB/RIF and Xpert MTB/RIF Ultra assays, our preference was to design Xpert MTB/XDR so that it had an analytical sensitivity at least matching that of the Xpert MTB/RIF assay, keeping in mind that the analytical sensitivity of Xpert MTB/RIF assay is roughly equivalent to the analytic sensitivity of the *rpoB* component of the Xpert MTB/RIF Ultra assay (36). The prototype cartridge LoD was 300 CFU/mL, which was in the range of, but not as good as, the Xpert MTB/RIF assay (130 CFU/mL). Our new Xpert MTB/XDR assay showed a comparable LoD to Xpert MTB/RIF for *M. tuberculosis* detection. We confirmed the reliability of our LoD estimation with multiple cartridge lots and analytical studies performed at two laboratories. We did not perform any head-to-head comparisons between Xpert MTB/XDR and Xpert MTB/RIF Ultra since we expect that Xpert MTB/XDR will perform well with samples that test positive by Xpert MTB/RIF Ultra as long as the Xpert MTB/RIF Ultra does not produce a “Trace” call with “Indeterminate” rifampin resistance results due to low bacillary load.

We performed a limited clinical study on a panel of frozen sputum and clinical isolates from three different continents representing a wide range of clinically relevant mutations. The clinical isolates and sputum samples represent a considerable geographical variation, and thus enabled us to assess the performance of the assay as a reflex test on both TB positive sputum samples as well as *M. tuberculosis* clinical isolates. The performance of the assay when compared to P-DST generated very high sensitivity and specificity values except for ETH, which showed a sensitivity of 65.4%. The assay showed 100% specificity and 94-100% sensitivity in detecting WT and mutant sequence types for all other drug targets. The low sensitivity for ETH when compared to either reference standards can be explained by the fact that this assay targets only mutations in the *inhA* promoter, among the several other possible gene mutations, which may be associated with ETH resistance (16). In our study sample group, there were several ETH resistance-associated mutations in the *ethA* and the *ethA* upstream region, which are not targeted by our assay, which accounted for the low sensitivity of the assay in detecting ETH resistance. Detection of ETH resistance was not an original aim of the assay and was included later, since ETH resistance has been reported to show significant association with mutations in *inhA* promoter that are also associated with low level INH resistance as tested by our assay(35) (41). We can expect that that the assay will show similar performance when tested in a larger multi-centric clinical study.

The Xpert MTB/XDR assay is intended to be used as a reflex test for a specimen that is determined to be *M. tuberculosis* positive and to serve as an aid in the diagnosis the main types of resistance that exist in M/XDR TB when used in conjunction with clinical and other laboratory findings. To address the global MDR-TB crisis and expedite diagnosis, WHO has determined that expanding rapid testing and the detection of drug-resistant TB is a top priority (42) and recently endorsed a 6-9 month shorter treatment regimen, replacing conventional 18-24 month regimens (43). Access to fast, sensitive, and safer genotypic assays like Xpert MTB/RIF Ultra and Xpert MTB/XDR, which detect resistance by identifying mutations known to confer resistance to the first- and second-line drugs in a majority of clinical strains, will minimize the biohazard and reduce sample preparation to a few manual steps that are more amenable to use at the point of care. When used as a reflex assay in conjunction with Xpert MTB/RIF or Xpert MTB/RIF Ultra, the Xpert MTB/XDR assay can expand TB and drug resistance detection to medically underserved populations.

## Funding

Research reported in this publication was supported by the National Institute of Allergy and Infectious Diseases of the National Institutes of Health under Award Number R01AI111397 and a grant from the Foundation for Innovative New Diagnostics. Research support was also provided by Cepheid. Cepheid collaborated in assay design, analytical study design and performance, while FIND was involved in the clinical study planning, design, and providing samples. The NIH had no role in study design, planning, or manuscript preparation. The content of this article is solely the responsibility of the authors and does not necessarily represent the official views of the National Institutes of Health.

## Conflicts of interest

D.A. receives income from license payments from Cepheid to Rutgers University. D.A. also reports receiving research contracts and support from Cepheid. D.A. and S.C. report the filing of patents for primers and probes for detecting drug resistance in *M. tuberculosis*. R.L.G., D.L., S.R., N.V., R.K., D.P. and S.C. are employed by Cepheid.

